# Incorporating selfing to purge deleterious alleles in a cassava genomic selection program

**DOI:** 10.1101/2020.04.04.025841

**Authors:** Mohamed Somo, Jean-Luc Jannink

## Abstract

Cassava has been found to carry high levels of recessive deleterious mutations and it is known to suffer from inbreeding depression. Breeders therefore consider specific approaches to decrease cassava’s genetic load. Using self fertilization to unmask deleterious recessive alleles and therefore accelerate their purging is one possibility. Before implementation of this approach we sought to understand better its consequences through simulation. Founder populations with high directional dominance were simulated using a natural selection forward simulator. The founder population was then subjected to five generations of genomic selection in schemes that did or did not include a generation of phenotypic selection on selfed progeny. We found that genomic selection was less effective under the directional dominance model than under the additive models that have commonly been used in simulations. While selection did increase favorable allele frequencies, increased inbreeding during selection caused decreased gain in genotypic values under the directional dominance. While purging selection on selfed individuals was effective in the first breeding cycle, it was not effective in later cycles, an effect we attributed to the fact that the generation of selfing decreased the relatedness of the genomic prediction training population from selection candidates. That decreased relatedness caused genomic prediction accuracy to be lower in schemes incorporating selfing. We found that selection on individuals partially inbred by one generation of selfing did increase mean genetic value of the partially inbred population, but that this gain was accompanied by a relatively small increase in favorable allele frequencies such that improvement in the outbred population was lower than might have been intuited.

## Introduction

Over the evolution of wild ancestors of crop plants, during domestication, and during breeding, crop populations can accumulate deleterious alleles (Valluru et al., 2019; Ramu et al., 2017). This problem may be more severe in clonally-propagated crops, such as cassava, because of the many propagation cycles they undergo between meiotic recombination events. The propagation cycles mean more rounds of DNA replication leading to opportunities for mutation whereas recombination facilitates purging by decreasing the association of favorable and deleterious alleles across loci. Cassava has been found to carry high levels of deleterious mutations, distributed throughout the genome (Ramu et al., 2017). Ramu et al. (2017) further found that over the course of modern breeding, deleterious alleles have become more common, but that performance among cultivars has been maintained by the masking of these alleles in heterozygous state. Given the recessive mode of action of these alleles, bringing them to homozygous state could be a way to detect and purge them.

Selfing and subsequent inbreeding depression leading to fitness loss can cause deleterious recessive allele purging under natural selection (Charlesworth &Willis, 2009; Lande et al., 1994). Although rounds of selfing can help purge recessive deleterious alleles, it is less efficient against mildly deleterious alleles. This means that while selfing could help eliminate strongly deleterious alleles it might also accelerate the fixation of mildly deleterious alleles (Boakes & Wang, 2005).

Selfing in cassava is associated with inbreeding depression and loss of fitness (Rojas et al., 2009; Nuwamanya et al., 2011; Ceballos et al., 2004). The expression of recessive alleles has been used for gene discovery. For example, genotypes carrying waxy genes associated with starch were S1 derived from AM206-5 clone (Ceballos et al., 2007). Similarly, Prochnik et al. (2012) used the predominantly homozygous AM560-2 cassava clone (S3 derived from MCOL-1505) for genome sequencing. Studies have reported varied inbreeding depression on different cassava traits. Both Rojas et al. (2009) and Nuwamanya et al. (2011) examined the effect of a single round of selfing on fresh root yield and found 60% reduction in yield. Kawuki et al. (2011) reported a 36% decrease of dry matter following a cycle of selfing. Finally, de Freitas et al. (2016) found that inbreeding depression varied widely between clones, ranging from 2% to 55% for fresh root yield, 0% to 9% for height, and 0% to 2% for dry matter content.

With GS breeding schemes being adopted in cassava breeding (Wolfe et al., 2017), there is a higher risk of inbreeding because cycles of selection are more frequent and each cycle entails a bottleneck (Ozimati et al., 2019). Ozimati et al. (2019) compared inbreeding between C0 and C1 populations using the average of diagonal elements of a kinship matrix, as a measure of the inbreeding coefficient. While they found less inbreeding in the C1 than in the C0 population, they attributed that to a purposeful crossing design that prevented genomically similar individuals from being crossed. Other GS studies, as well as simulation, have found a potential for rapid loss of diversity (Rutkoski et al., 2015; Jannink, 2010).

Empirically testing the impact of selfing cycles for cassava with a relatively long life cycle would be slow and expensive so that breeders are reluctant to explore selfing as a component of the GS scheme. To our knowledge, no GS breeding studies with selfing have been conducted to estimate the effect that selfing might have on gain from selection. In addition to empirical tests, stochastic simulation can be used to test breeding schemes and gain insight into the impact of specific changes such as incorporating selfing. In this study our objectives were to (1) compare breeding schemes with and without selfing in scenarios with different training population sizes and selection intensities under directional dominance and additive modes of gene action, (2) identify mechanisms affecting the observed responses to selection in terms of changes in the frequency of favorable alleles and deleterious homozygote genotypes.

## Materials and methods

### Generation of founder haplotypes using forward simulation

Because the cassava genome has been found to harbor many deleterious recessive mutations (Ramu et al., 2017), we chose to start breeding schemes from founder haplotypes with evolved directional dominance, induced both by the mode of gene action and historical natural selection. We generated starting populations using the forward simulator SLiM (Haller & Messer, 2017). SLiM is a stochastic simulator enabling flexible simulation at the base pair level. Our simulations involved evolutionarily constrained segments that we call here “genes” though superficial evaluation shows that these segments resemble the genes of molecular geneticists poorly. Likewise, “base pairs” in our genes underwent stochastic mutations with different probabilities and severity of deleteriousness. Our objective was to generate haplotypes with directional dominance including some level of complexity in their gametic phase disequilibrium and allele frequency spectrum. We simulated genomes with five chromosomes of 30 Mbp and 90 cM length each. These chromosomes were interspersed with genes every 100,000 bp (300 genes per chromosome). Genes were 20,000 bp in length, with characteristic mutation effects and mode of action dependent on position within the gene (Table 1). The simulation gave genes rather long regions subject to mutations with milder effects where the deleterious allele was only partially recessive, and more constrained regions where mutations had stronger effects, were more frequently deleterious, and where the deleterious alleles were more completely recessive. The overall mutation rate was 1e-7 per base pair per generation. The forward simulation was run for 12,000 generations with a census population size of 500.

We recognize that the cassava genome is much different from the genome that we simulated, with 770 million base pairs of sequence, 18 chromosomes and 33,033 annotated genes. In recombination space, the genome spans 2412 cM (International Cassava Genetic Map Consortium, 2015). We nevertheless believe that the genome we simulated can provide instructive insights.

**Table 1.** Mutation characteristics as a function of region in genes. The selection coefficient *s* gives the fitness of the homozygote mutant (*W* = 1 + *s*) relative to the non-mutant (*W* = 1). The dominance coefficient h gives the fitness of the heterozygote (*W* = 1 + *hs*). The Deleterious Frequency indicates the percentage of mutants in the region that are deleterious.

| Base pair position of region | Deleterious Frequency (%) | Deleterious |  | Favorable |  |
| --- | --- | --- | --- | --- | --- |
| | | $s$ | $h$ | $s$ | $h$ |
| 1 – 10,000 | 90 | -0.004 | 0.20 | 0.004 | 0.80 |
| 10,001 – 14,000 | 95 | -0.008 | 0.10 | 0.004 | 0.80 |
| 14,001 – 16,000 | 98 | -0.016 | 0.05 | 0.008 | 0.90 |
| Two region types alternating every 200bp | 98 | -0.030 | 0.02 | 0.008 | 0.90 |
| 16,001 – 20,000 | 95 | -0.008 | 0.10 | 0.004 | 0.80 |

### Breeding scheme simulation

The BreedingSchemeLanguage (BSL) R Package (Yabe et al., 2017) was used to simulate breeding schemes. We assumed there were no new mutations during the breeding program. Simulations in SLiM parameterized mutations in terms of their effects on fitness. The BSL environment, however, parameterized mutations in terms of their effects on a quantitative trait. In the BSL, then, the effect of the trait was proportional to the *s* value shown in Table 1, and its degree of dominance was the same as in SLiM, such that the favorable (high-valued) allele was always dominant over the deleterious allele. The initial genetic variance was set to 1. The founders were considered to have been well phenotyped, thus initial error variance was set to 1, leading to heritability 0.5 for founders. Error variances associated with subsequent evaluations at seedling (as is common in cassava, we refer to any plant grown from botanical seed as a “seedling” regardless of how old and big it gets), clonal evaluation trial (CET), and preliminary yield trial (PYT) plot types for model updating were set to 16, 9, and 4, respectively. The error variances for different evaluation types were held constant in each cycle of the breeding schemes.

We implemented a five-cycle scheme after initial evaluation and selection of founder populations. The standard scheme involving selfing was as follows. Founders’ genomic estimated breeding values (GEBVs) were estimated using ridge regression (Endelman, 2011) and sixty individuals with high GEBVs were selected as parents for the next generation. Those parents were selfed, creating 10 S1 progeny per parent. The resulting 600 partial inbreds were evaluated as seedlings. The five progenies with highest values were chosen from each of the 60 families for the crossing nursery. To generate 1,000 outbred progenies, the 300 individuals (60 selected parents x 5 S_1_ per parent) were randomly mated, excluding within-family matings. Outbred progenies were genotyped and their GEBVs estimated using all prior phenotypic data, again selecting the best 60 individuals. This scheme was repeated in subsequent cycles.

The genomic prediction training population started with the founders (cycle C0). After the creation of C3 progeny (but before they were selected), the training population was updated with phenotypic records of 440 non-parental cycle C1 individuals and the 60 C1 parents of C2 progeny using CET plots. After the creation of C4 progeny, the training population was updated with phenotypic records of 440 non-parental cycle C2 individuals and the 60 C2 parents of C3 progeny using CET plots as well as 190 non-parental C1 individuals and the 60 C1 parents of C2 progeny using PYT plots. The number of C5 progeny (the final generation) was set to 200. All breeding schemes were replicated 24 times.

### Simulation scenarios

In addition to the standard scheme with selfing, we simulated a no selfing scheme. There, the 60 selected parents were intermated at random leading directly to the next generation. We considered schemes with or without a selfing generation to have the same overall breeding cycle time. We also simulated a “selfing-plus” (Self+) scheme used to update the training population more rapidly. In the Self+ scheme, the phenotypic data from evaluating S1 progeny of selected parents were incorporated back into the training population by attributing to the parent the mean value of its ten S1 progeny. Standard schemes involved 600 founders as the initial training population. We also tested founder population sizes of 200 and 1500. Standard schemes involved 1,200 markers per chromosome. We also tested schemes with 3,600 markers per chromosome. Standard schemes involved a selection intensity of 10% among founders and 6% thereafter. We also tested selection intensities of 5% among founders and 3% thereafter. Finally, we simulated the standard schemes with an additive gene model to contrast with the directional dominance gene model.

### Analysis of simulation results

We tracked change in genotypic value across breeding cycles. The QTL could have effects of four different sizes (Table 1). We tracked mean favorable allele frequency across all loci within an effect class, so that “allele frequency” was actually calculated over many loci. We also tracked a composite favorable allele frequency as a weighted mean, with weights being the effect of the locus class (i.e., weights of 4, 8, 16, and 30, Table 1). Similarly, we tracked homozygote deleterious genotype frequency for which the frequency is over individuals rather than over gametes. We measured the impact of selection on inbreeding depression by calculating the loss of genetic value due to one generation of self-fertilization in the founder population as compared to the Cycle 5 population. For all of these measures, we performed simple ANOVA or t-tests for relevant comparisons across cycles or across simulation scenarios to determine significance of differences.

We called selection on GEBVs of outcrossed individuals as the “main selection” and on selfed S1 individuals as “purging selection.” Populations in schemes with no selfing were subject only to main selection while those with selfing underwent main selection and purging selection. Comparison of gain due to main and purging selection were done for additive and dominance models for the Self, NoSelf, and Self+ schemes, and across the three founder population sizes (200, 600, and 1500).

### Data availability

No empirical data were used or generated. The simulation and analysis scripts are deposited on github at https://github.com/jeanlucj/PurgingBySelfingSimulations.

## Results

### Genomic selection under directional dominance

We observed significant differences among simulated breeding schemes (Fig. 1). The Self and Self+ schemes achieved higher means than the NoSelf scheme. Qualitatively, the responses of all schemes followed a similar pattern: rapid gain in the first cycle, diminished gain in the second cycle, loss in the third cycle, and recovery in the fourth and fifth cycles (Fig. 1). This pattern meant that gain in genetic mean from generations 1 to 5 was not significant. Genomic prediction models lose accuracy after selection (e.g., Muir, 2007). To determine if the pattern observed here was simply caused by that mechanism or was caused by the mode of gene action, we simulated selection under additive gene action (Fig. 2). Under additivity, the pattern of response was very different, with steady gain observed across different founder population sizes (Fig. 2). Note that gain under the additive model was three to four times greater than under the directional dominance model, despite all schemes starting from populations with the same genetic variance. Variation in gain among replicate simulations was also lower under the additive than the dominance gene models, as reflected by the smaller standard errors of population means (Fig. 2). To better understand the response under directional dominance, we examined allele and genotype frequencies across the four classes of loci in the simulations. We wanted to distinguish two hypotheses. First, loss of accuracy of the prediction model would be expected to occur more quickly under directional dominance than additivity (Duenk et al., 2019). So, despite little loss of accuracy under the additive model, the question of a loss of accuracy for breeding value prediction under dominance was still relevant. Second, under directional dominance the genotypic value depends strongly on the frequency of the deleterious allele homozygote genotype frequency. Consequently, decreased response could be due to increased frequency of that genotype. Favorable allele and deleterious homozygote genotype frequencies supported the second hypothesis (Fig. 3). In particular, the breeding values of individuals was a function of the favorable allele frequencies (Falconer & Mackay, 1996), which trended upward for all classes of loci (Fig. 3), though certainly more so for the larger effect alleles. Favorable allele frequency changes were small, as is to be expected under an infinitesimal model. The trend upward shows the effectiveness of genomic selection to increase breeding values, even under directional dominance.

**Fig. 1.**
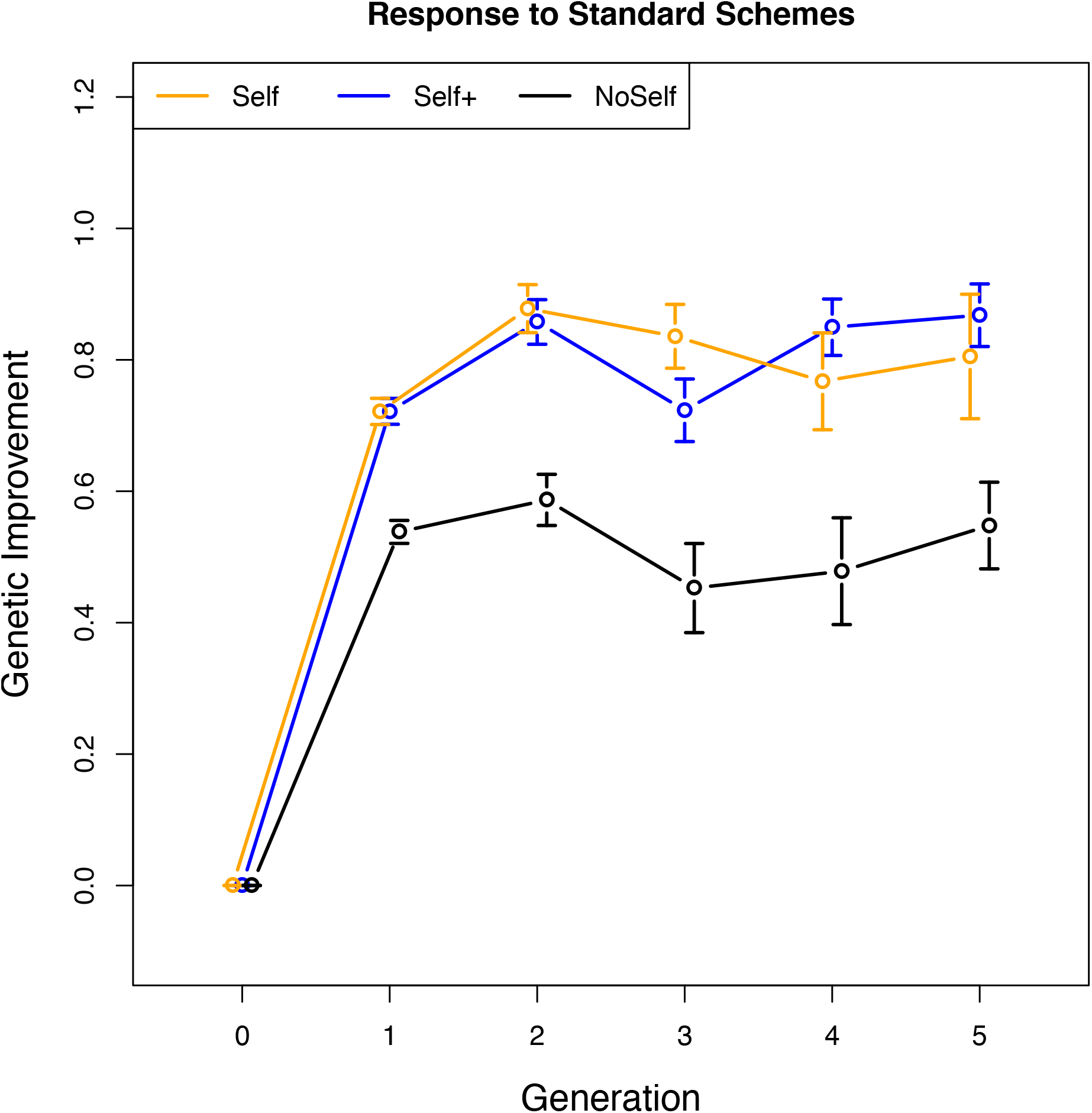
Gain from selection using two schemes with selfing (Self and Self+) compared to one without selfing (NoSelf) over five generations. The number of founders was 600.

**Fig. 2.**
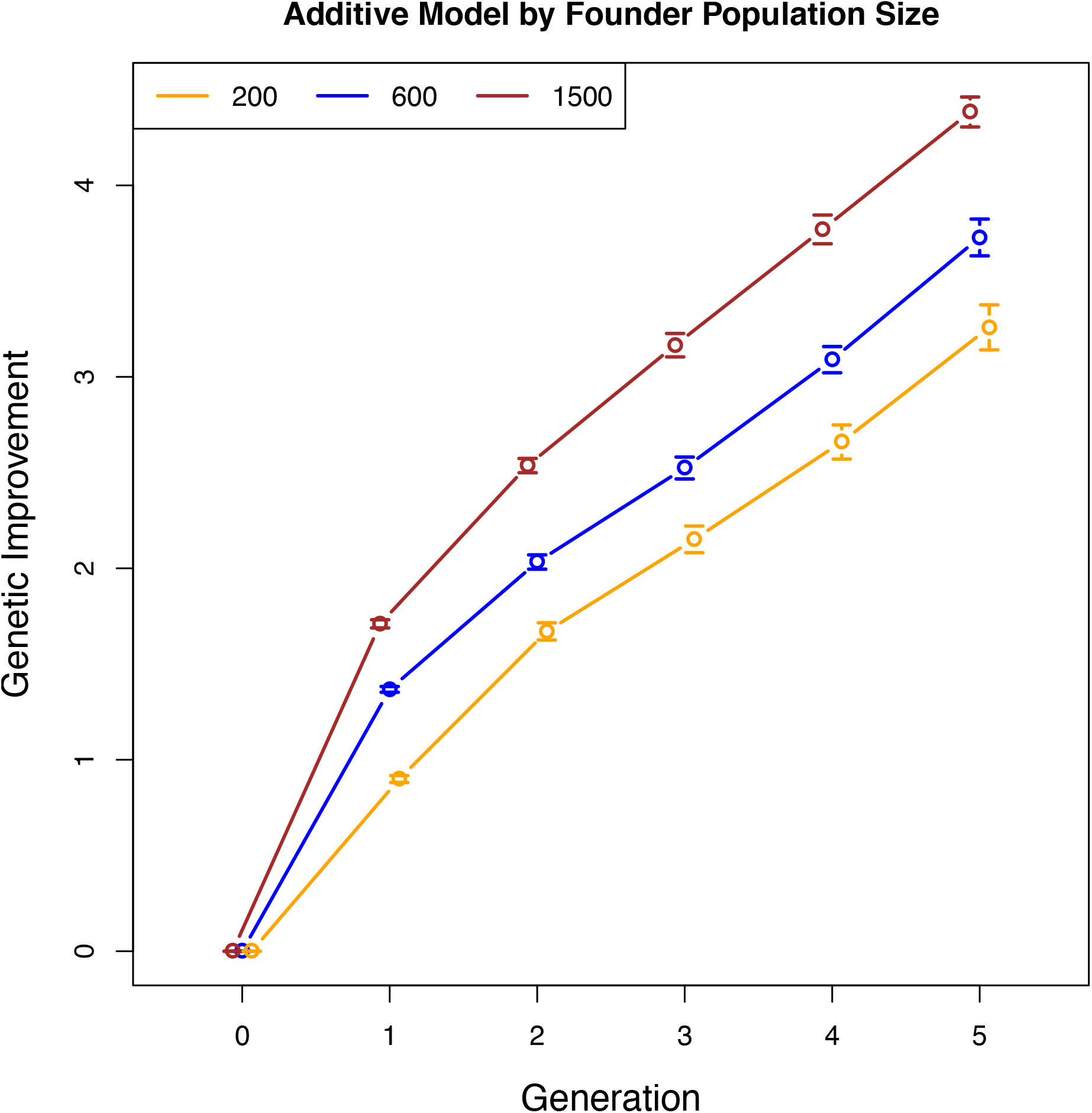
Gain from the Self breeding scheme under an additive gene action model, assuming different founder population sizes. Other than mode of gene action, all parameters were the same as for the breeding schemes shown in Fig. 1.

**Fig. 3.**
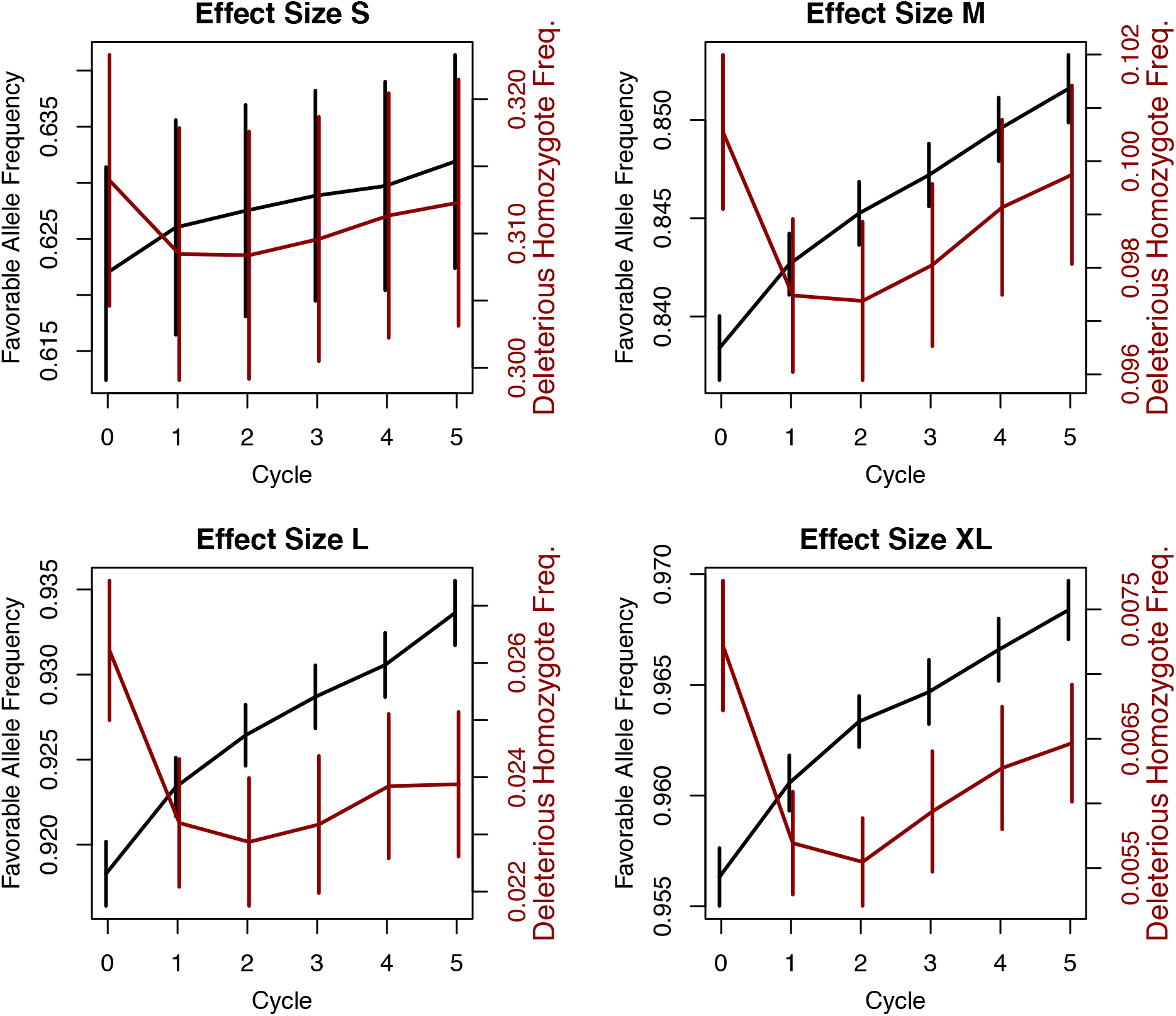
Favorable allele frequencies (black, left scale) and deleterious allele homozygote frequencies (red, right scale, note differences in scales) over five cycles of selection in the standard setting. Effect sizes of 0.04, 0.08, 0.16, and 0.30 labeled S, M, L, and XL, respectively. Error bars are +/- one standard error across the 24 repeated simulations.

Apparently paradoxically, however, the frequency of the deleterious allele homozygote genotype also increased. The trend for increase of favorable allele frequency with concomitant increase in the frequency of deleterious allele homozygotes was observed across all allele effect sizes. As a possible explanation for these opposing trends, we highlight the change in effective population size between the founders (*N_e_* of about 500) and during the breeding cycles (*N_e_* of about 60). That change meant that even as favorable alleles increased in frequency, variation in allele frequency across loci also occurred, leading to fixation at some loci. The hypothesis of fixation leading to higher frequencies of deleterious homozygote genotypes was supported by response to selection under high selection intensity (Fig. 4) and given a small founder population size (Fig. 5). In the case of high selection intensity, the number of individuals selected was lower, leading to more severe bottlenecks. Comparing Fig. 4 to Fig. 1, the higher selection intensity leads to greater gain in the first selection event than the standard scenario, but in subsequent generations the populations crash, leading to negative responses to selection (Fig. 4). Interestingly, the Self+ scheme, with its more rapid (though limited) updating of the training population to some extent rescues the populations from this crash.

**Fig. 4.**
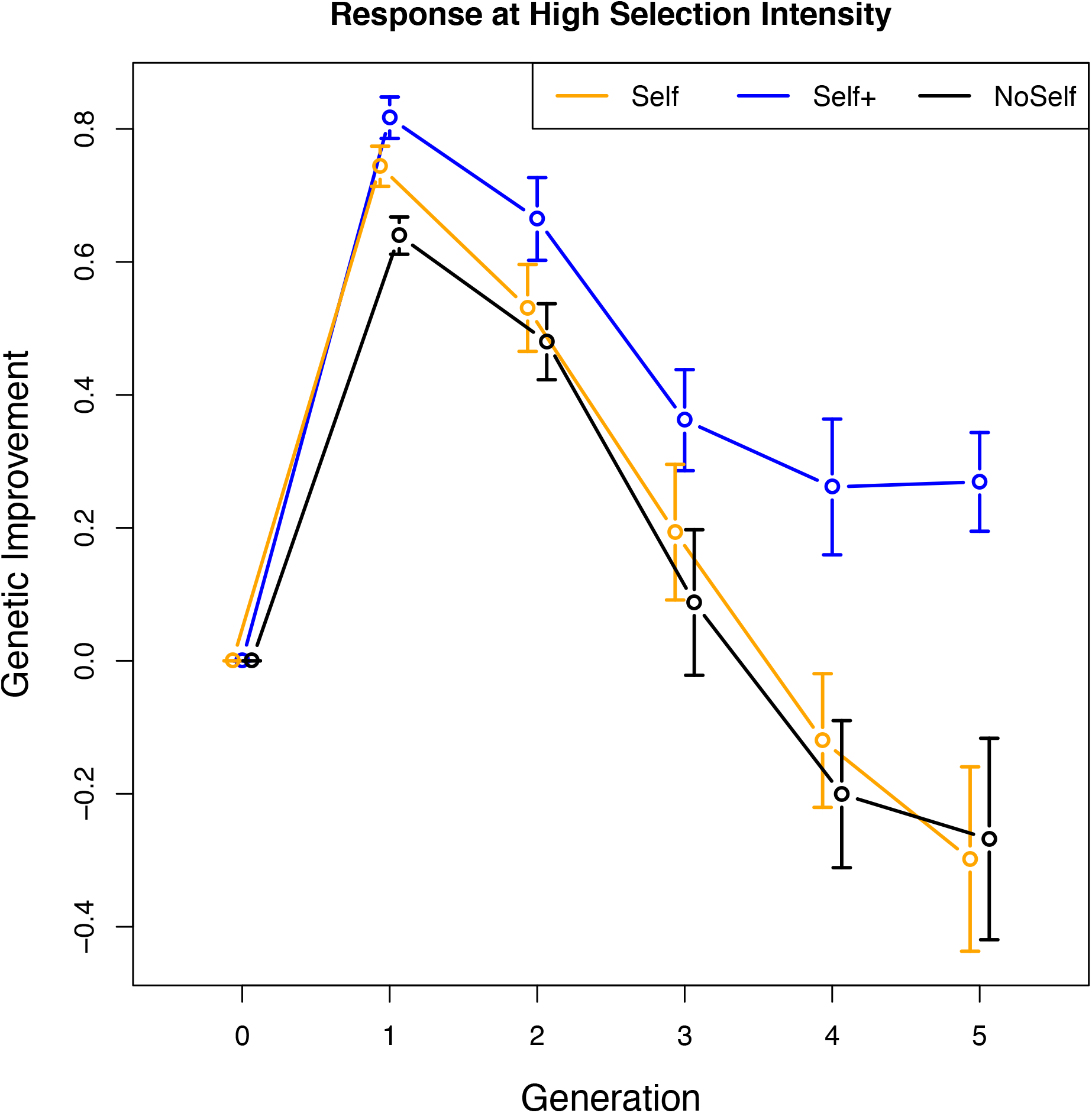
Gain from selection using two schemes with selfing (Self and Self+) compared to one without selfing (NoSelf) over five generations. All parameters were the same as for the breeding schemes shown in Fig. 1 with the exception that 30 parents were selected in each cycle as opposed to 60.

**Fig. 5.**
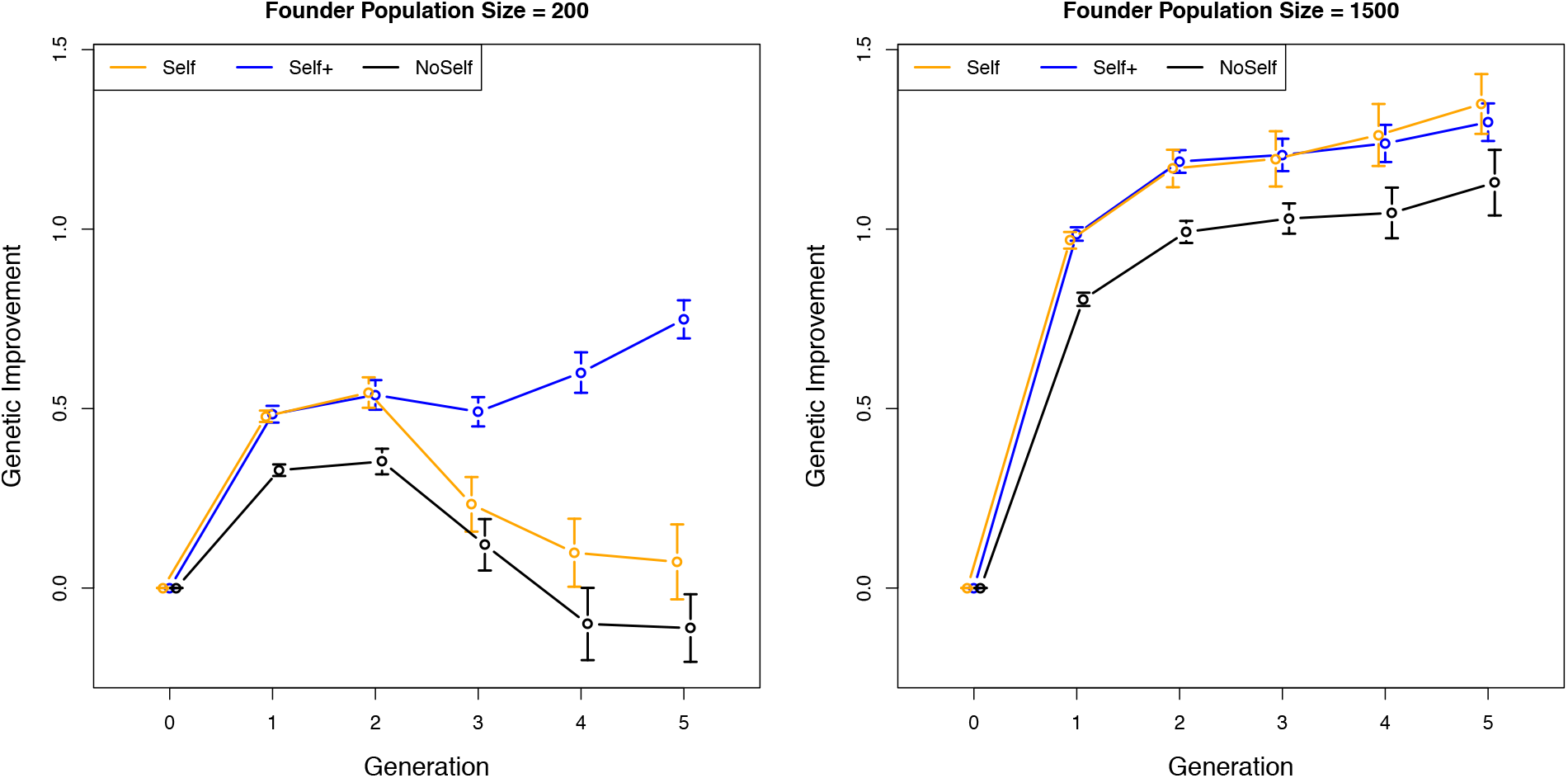
Gain from selection using two schemes with selfing (Self and Self+) compared to one without selfing (NoSelf) over five generations. All parameters were the same as for the breeding schemes shown in Fig. 1 with the exception that founder population sizes were 200 and 1500 as opposed to 600.

In the case of small founder population size, it is known that with a smaller training population, genomic selection causes higher co-selection of relatives and therefore more rapid loss of diversity / inbreeding (Jannink et al., 2010). With the small founder population (and therefore smaller training population) we again saw a population crash and negative responses to selection (Fig. 5). The Self+ scheme again rescued these populations from a total crash. In contrast, with the large founder population of 1500 individuals, there was never a decline in the population mean, and the Self and Self+ schemes perform equally well (Fig. 5). As for the standard breeding scheme, the mechanism generating dramatic drops in genotypic value had to do with the accumulation of deleterious homozygotes in the population. For both the high intensity selection and the small founder population breeding scenarios, the Self scheme had higher rates of deleterious homozygote accumulation than the Self+ scheme (Supp. Fig. 1), even though the increase in the favorable allele frequency was similar across schemes. For the large founder population breeding scenario, Self and Self+ schemes did not differ, with low increase in deleterious homozygote frequency and greater increase in favorable allele frequency.

### Selection efficiency after the first cycle with and without selfing

Another observation from the standard scheme was that all of the increased gain in the Self and Self+ schemes relative to the NoSelf scheme occurred in the first two cycles of selection (Fig. 1). Thereafter, the gains of the Self and Self+ schemes were not significantly different from that of the NoSelf scheme. The selfing schemes benefit from two selection events leading to genetic gain, the main event from genomic selection on outcrossed individuals and the purging event on selfed individuals. For the selfing schemes not to do better than the NoSelf scheme requires that one or both of these events must decrease in effectiveness in the selfing schemes after the first two cycles of selection. In fact, only the main event decreased in effectiveness during selection events 3 and 4 (Fig. 6). We hypothesize that the reason for the drop in genomic prediction accuracy in the selfing but not the NoSelf scheme is that, by adding a generation between the training and selection candidate populations, the degree of relationship between those populations was lower in the selfing schemes than in the NoSelf scheme. The degree of relationship has a large impact on prediction accuracy (Clark et al., 2012).

**Supp. Fig. 1.**
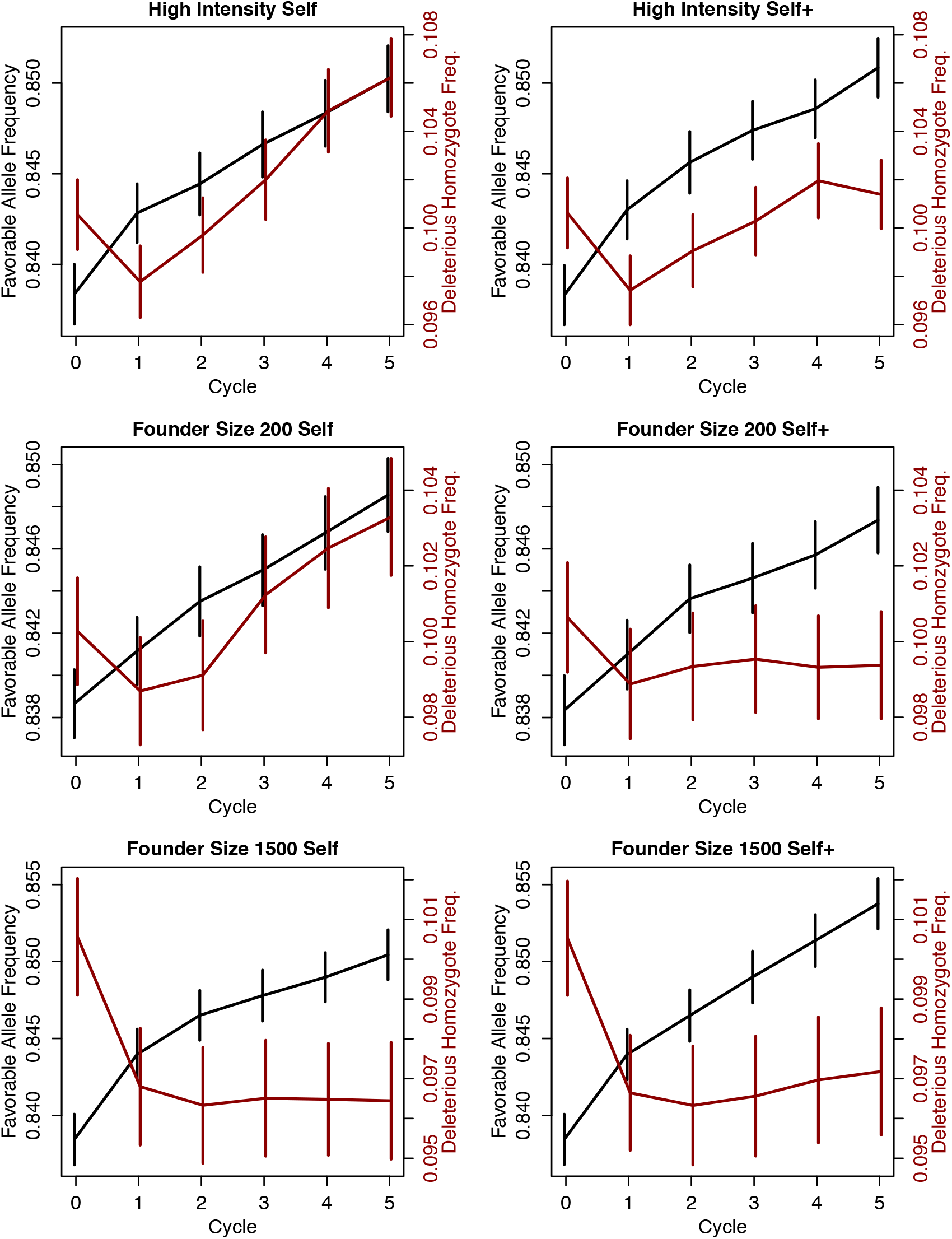
Weighted mean of favorable allele frequencies (black, left scale) and deleterious allele homozygote frequencies (red, right scale, note differences in scales) over five cycles of selection. Weights were 0.04, 0.08, 0.16, and 0.30 for the four locus classes, S, M, L, and XL, respectively. Error bars are +/- one standard error across the 24 repeated simulations. Left and right columns show the Self and Self+ breeding schemes, respectively. Top, middle, and bottom rows show founder population size of 600 with high selection intensity, and founder population size of 200 and 1500 with standard selection intensity, respectively.

**Fig. 6.**
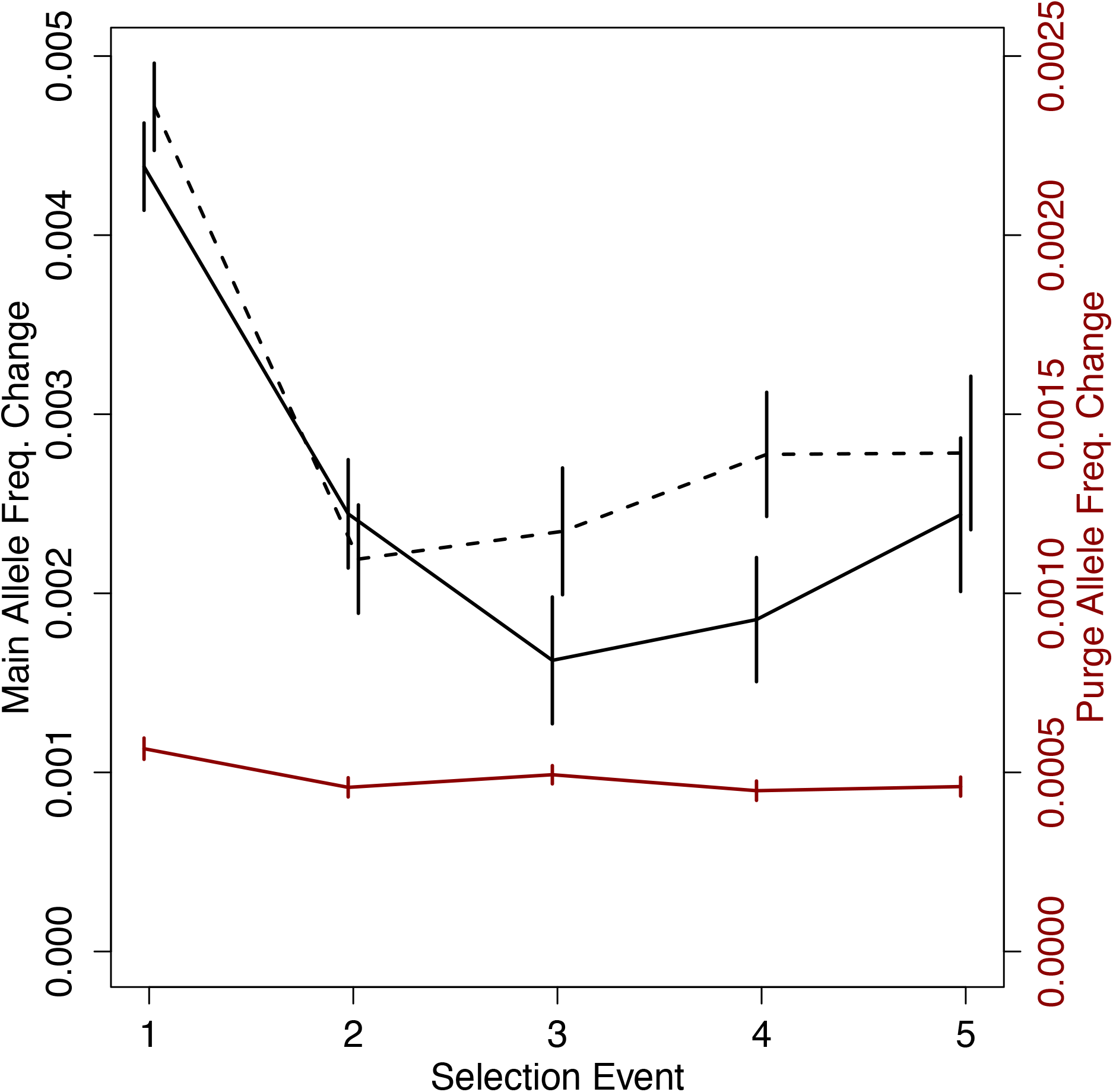
Weighted mean of favorable allele frequencies during the main (black, left scale) and purging (red, right scale, note differences in scales) selection events in the standard selection scheme. Weights were 4, 8, 16, and 30 for the four locus classes, S, M, L, and XL, respectively. Error bars are +/- one standard error across the 24 repeated simulations.

**Fig. 7.**
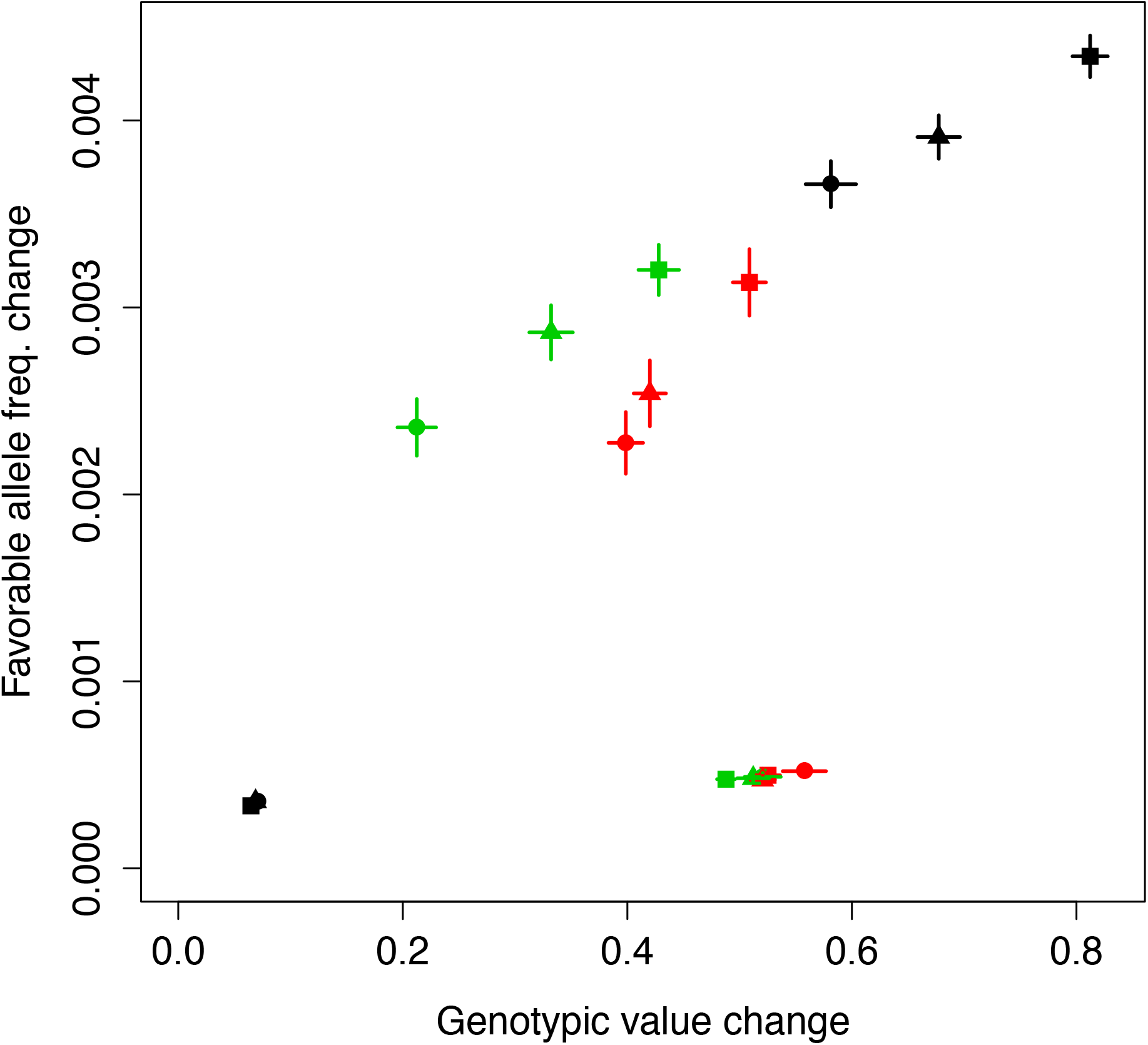
Comparison of genotypic value change versus weighted favorable allele frequency change caused by selection events. Black, red, and green symbols are, respectively, for additive gene action, and dominance with Self and Self+ selection. Round, triangle, and square symbols are, respectively, for founder populations sizes of 200, 600, and 1500. Favorable allele frequency changes under 0.001 occurred during purge selection, while changes above 0.002 occurred during main selection. Whiskers show one standard error above and below the observed value. An absence of whiskers indicates a standard error smaller than the size of the symbol.

### Inbreeding depression

We compared inbreeding depression before versus after the five cycles of selection by comparing the loss of genetic value caused by one generation of self-fertilization, either on the founder population or on the Cycle 5 population. As expected, inbreeding among founders for all breeding scenarios were not significantly different (given that no breeding treatment had yet been applied). One generation of selfing on a founder population caused a mean decrease of 5.63 in the genetic value, with a standard deviation of 0.38 across replications. After five cycles of selection, there were significant differences among selection schemes, though, surprisingly, the difference was that inbreeding depression was higher under the Self+ scheme while the Self and NoSelf schemes showed similar but lower inbreeding depression (Table 2). Equally surprisingly, the number of founders was not significantly related to decrease in inbreeding depression (Table 2). We hypothesize that the level of inbreeding depression was due not only to the extent of deleterious allele purging, but also to the level of inbreeding previously attained in the population. Thus, there was greater inbreeding depression under the Self+ than the Self scheme not because it was less effective at purging but because the Self+ population was less inbred than the Self population (Fig. 3, Supp. Fig. 1). This hypothesis could also explain why inbreeding depression was as great given a 1500 versus a 200 founder population: the former was more effective at purging but also was less inbred than the latter (Supp. Fig. 1).

**Table 2.**
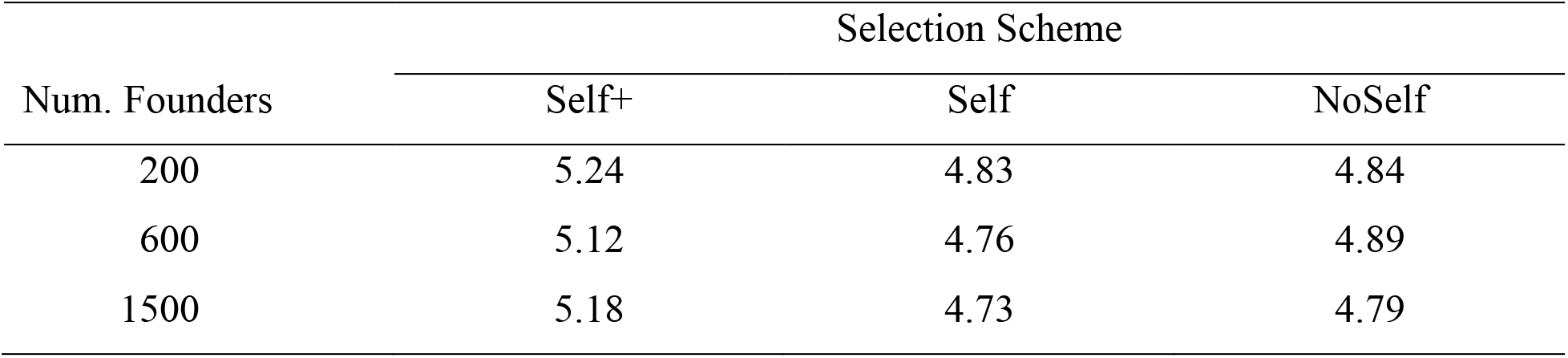
Inbreeding depression from one generation of selfing after five cycles of selection for different breeding schemes. Means across 24 replications are reported and have a standard error of the mean of 0.08.

## Discussion

This study investigated the effect of purging recessive deleterious alleles by selection on selfed individuals during five generations of genomic-assisted breeding. Perhaps most obviously, we showed that response under directional dominance differs dramatically from that under an additive model (contrast Fig. 1 and 2). By analyzing changes in favorable allele frequency and deleterious homozygote genotype frequency, we showed that the sudden decline in response under directional dominance was caused by drift fixing deleterious alleles at some loci, even as the favorable allele frequency, averaged over all loci, increased (Fig. 3). These opposing effects led to stagnating or declining genotypic values. Support for this interpretation also came from simulation scenarios with higher selection intensity (hence more drift, Fig. 4) and smaller training population size (hence more co-selection of relatives, Fig. 5).

On the strict question of the value of selecting among selfed progeny, our results did not lead to an unambiguous answer. First, selection among selfed progeny was effective in the sense that it led to a large shift in genotypic value among those progeny (Fig. 7). This shift, however, did not lead to a large shift in the favorable allele frequency (Fig. 7). The change in favorable allele frequency during purging selection was only about one fourth of that during main selection (Fig. 6). The change in genotypic value was of course affected by the simulation parameters we assumed: relatively high error variance among seedlings and low selection intensity during purging selection. Empirical estimates of error variance or broad sense heritability for yield traits in cassava seedlings have not been reported (e.g., Ozimati et al. 2019). Such estimates would be difficult to obtain given that all seedlings are genetically unique and cannot be replicated. An estimate of the residual from additive effects, which would be straightforward to estimate, would include a component of non-additive genetic variation. In the presence of dominance, that component could be considerable. Our choice of a high error variance on the seedlings was also influenced by the fact that cassava storage root properties differ between seedling and clonal plants (Ceballos et al., 2004), so that lack of genetic correlation between seedling and clonal performance would also contribute to the error of seedling measures to predict clonal performance. Thus, it seemed reasonable to assume that the error variance on a single plant would be substantially greater than on a clonal evaluation performed on multiple plants. We justified the low selection intensity (five plants chosen out of ten grown) only as a matter of labor and cost savings. Obtaining many selfed progeny and planting them all for a large number of selected parents would be logistically challenging.

While the change in allele frequency from purging selection was low, the increase in genetic value among S1 progeny from that selection event was similar to that observed in the mean selection (Fig. 7). Under additive gene action (black symbols in Fig. 7), there was a direct relationship between the genotypic value and the favorable allele frequency changes, regardless of main versus purge selection. Under additivity, however, purging selection caused a smaller genotypic value change and thus a smaller favorable allele frequency change (Fig. 7). Under directional dominance gene action, while purging selection caused greater genotypic value change than main selection, it caused little favorable allele frequency change (Fig. 7). Empirical selection studies using selfing have sometimes also observed disappointing gains (Wardyn et al., 2009). Under purging selection, individuals’ genotypic values depend on how many deleterious homozygote loci they carry. But there is not a perfect correlation between the number of homozygous loci and the total number of deleterious alleles, which may also be in the heterozygous state. In other words, genotypic value offers an uncertain guide to favorable allele content, such that the strong shift of mean genotypic value under purging did not correspond to a strong shift in favorable allele frequency. We also found that while selection on selfed individuals did lead to a reliable increment in breeding value in each cycle, it also caused the genomic prediction model accuracy to decrease (Fig. 6). We assume that the effect was due only to the addition of a generation between the training population and the selection candidates, making them more distantly related (Clark et al. 2012). That effect erased the benefit of the purging selection. It is unclear whether the accuracy drop would have persisted over further cycles of selection, as the training population began to be updated (Fig. 6). We opted to simulate only five cycles of selection because five cycles actually represent a long period of time, a minimum of ten years. In that period of time, it is unlikely that any single breeding scheme would persist, given technological and breeding priority changes. The value of simulating longer time periods is therefore unclear. Finally, we note that we did not impose any penalty on the selfing schemes for the amount of time that adding the generation of selfing would require. That time would decrease the number of breeding cycles possible per unit time, an effect that has strong negative repercussions (Cobb et al., 2019).

## Conclusions

Cassava carries a high genetic load (Ramu et al., 2017) and is known to suffer from inbreeding depression (Ceballos et al., 2004). These observations suggest attempting to purge deleterious load by selection on partially inbred individuals could be worthwhile. We simulated one approach to implement purging selection in the context of a breeding program using genomic selection. We observed a favorable initial response (cycles 1 and 2) to adding a generation of selfing to the breeding scheme, but this benefit did not persist beyond those cycles. Over subsequent cycles (cycles 3 to 5), the increase in favorable allele frequency occurring during purging selection only compensated for the loss in accuracy a selfing cycle caused during the main genomic prediction cycle. Thus, despite added cost and overall breeding cycle length from including selfing, no net gain was observed relative to a scheme without selfing.

It is difficult to extrapolate from our results to what a cassava breeder may experience empirically because the results do depend both on questions of underlying genetic architecture and on breeding scheme details. Thus, it is difficult to make strong recommendations. We did find, however, a few somewhat non-intuitive results that we believe breeders should keep in mind as they consider breeding scheme modifications. First, we found that the breeding schemes increased favorable allele frequencies even though genotypic values were stagnant (Fig. 1 and Fig. 3). While we believe that this phenomenon can occur generally, it depended on a specific aspect of the simulation that will differ across real breeding programs. In particular, our simulated founders came from a population with an effective size of 500, whereas the standard breeding scheme had an upper bound to the effective population size of 60 due to the number of parents randomly mated in each cycle. This rapid downward change led to greater drift and therefore fixation of the deleterious allele at some loci. Thus, the lack of gain from the first to the fifth cycles (Fig. 1) in our simulation may not be generalizable to practicing breeding programs.

An observation that we believe will be generalizable was that introducing a generation of selfing into the scheme will have conflicting impacts: on the one hand, it will provide some genetic gain itself. On the other hand, it will distance selection candidates from the genomic prediction training population, thereby causing a decrease in accuracy. In our simulation, these conflicting impacts balanced out beyond the first selection cycle so that the selfing schemes generated about equal gain to the scheme without selfing (Fig. 1, 4, and 5). A related observation was that rapid additions to the training population in the Self+ scheme caused decreased inbreeding (Fig. 4, 5, and Supp. Fig. 1). We also think that this result is generalizable. We do not have an explanation as to why this effect did not lead to greater gains per cycle in the Self+ than the Self scheme. We do think the effect lead to greater potential for inbreeding depression at the end of the five selection cycles (Table 2), simply because the standing population was less inbred to begin with.

Finally, we observed that selection on partially inbred individuals led to a large gain in genotypic value among those individuals, but that that large gain was not reflected in a large gain in favorable allele frequency (Fig. 7). Under strong directional dominance, genotypic value may not be a good guide to breeding value. It is worth considering this effect more carefully. Changes in genotypic value are what breeders observe directly and it therefore guides their intuition. Here we show through simulation that selection on partially inbred individuals may not lead to as strong gain in their outcrossed progeny as we might intuit.

## Acknowledgments

The research conducted at Cornell University was part of NextGen cassava project funded by Bill & Melinda Gates foundation and UKaid (Grant 1048542). The research was conducted using the resources of the Cornell University Institute of Biotechnology Bioinformatics Facility (BioHPC). The authors gratefully acknowledge the support of BioHPC staff.

## Abbreviations

BSL: Breeding Scheme Language
CET: clonal evaluation trial
GEBV: genomic estimated breeding value
GS: genomic selection
PYT: preliminary yield trial

